# Cryptic proteins translated from deletion-containing viral genomes dramatically expand the influenza virus proteome

**DOI:** 10.1101/2023.12.12.570638

**Authors:** Jordan N Ranum, Mitchell P Ledwith, Fadi G Alnaji, Meghan Diefenbacher, Richard Orton, Elisabeth Sloan, Melissa Guereca, Elizabeth M Feltman, Katherine Smollett, Ana da Silva Filipe, Michaela Conley, Alistair B Russell, Christopher B Brooke, Edward Hutchinson, Andrew Mehle

**Affiliations:** Medical Microbiology and Immunology, University of Wisconsin-Madison, Madison WI 53706 USA; Department of Microbiology, University of Illinois, Urbana, IL 61801, USA; MRC-University of Glasgow Centre for Virus Research, Glasgow G61 1QH, UK; Division of Biological Sciences, University of California, San Diego, La Jolla, CA 92093 USA; Carl R. Woese Institute for Genomic Biology, University of Illinois, Urbana, IL 61801, USA

## Abstract

Productive infections by RNA viruses require faithful replication of the entire genome. Yet many RNA viruses also produce deletion-containing viral genomes (DelVGs), aberrant replication products with large internal deletions. DelVGs interfere with the replication of wild-type virus and their presence in patients is associated with better clinical outcomes as they. The DelVG RNA itself is hypothesized to confer this interfering activity. DelVGs antagonize replication by out-competing the full-length genome and triggering innate immune responses. Here, we identify an additionally inhibitory mechanism mediated by a new class of viral proteins encoded by DelVGs. We identified hundreds of cryptic viral proteins translated from DelVGs. These <u>D</u>elVG-encoded <u>pr</u>oteins (DPRs) include canonical viral proteins with large internal deletions, as well as proteins with novel C-termini translated from alternative reading frames. Many DPRs retain functional domains shared with their full-length counterparts, suggesting they may have activity during infection. Mechanistic studies of DPRs derived from the influenza virus protein PB2 showed that they poison replication of wild-type virus by acting as dominant-negative inhibitors of the viral polymerase. These findings reveal that DelVGs have a dual inhibitory mechanism, acting at both the RNA and protein level. They further show that DPRs have the potential to dramatically expand the functional proteomes of diverse RNA viruses.

## Introduction

RNA viruses face a race against the clock. Once infection starts, the virus must quickly replicate its genome and exit the cell before antiviral responses can mount an effective defense. This prioritizes speed, especially for acute infections like influenza virus. But this need for speed comes at a cost – fidelity is sacrificed as the genome is copied quickly by the viral RNA-dependent RNA polymerase (1). The speed and lack of proof reading by the polymerase results in the frequent introduction of point mutations into the viral genome. The process introduces approximately 10^−4^ to 10^−6^ substitutions per nucleotide per strand copied (2–4). These subtle changes allow rapid viral evolution, and are one of the underlying reasons why influenza vaccines must currently be reformulated each year (5).

RNA virus polymerases are also prone to larger replication errors. Instead of mutating a single nucleotide, these errors create large internal deletions in genome segments (6, 7). Most RNA viruses produce some version of deleted genomes, including important human pathogens like influenza virus, measles virus, SARS-CoV-2, and respiratory syncytial virus (8, 9). Influenza virus deletion-containing viral genomes (DelVGs, sometimes referred to as defective viral genomes, DVGs) are aberrant replication products containing a deletion of more than 10 nts while retaining the *cis*-acting sequences necessary for replication, transcription and packaging into virions (6, 10–13). Their smaller size gives them a competitive advantage during replication compared to their much larger wild-type (WT) counterparts (14). Indeed, the preferential replication of DelVGs is seen in high-throughput sequencing data where they cause they cause an under-representation of the deleted regions in the middle of viral genome segments (15). DelVGs originate from all eight influenza virus genome segments during infections in culture and in humans, but tend to be most abundant on the three largest genes that code for the viral polymerase subunits and possibly the next largest gene, hemagglutinin (13, 16–21). When DelVGs are packaged into virions, they create defective interfering particles (DIPs). Because DelVGs and the DIPs they form no longer encode essential viral proteins, they must parasitize WT viruses for their own replication (22). In so doing, they suppress replication of the WT virus. This interference phenomena is not restricted to influenza virus, and is a common feature of most RNA viruses (8).

DelVGs arise naturally in human and animal infections (18–21, 23). DelVG abundance is inversely correlated with the severity of influenza disease (24–26). As such, DelVGs and DIPs have been considered as alternative therapeutics, and recently used to disrupt SARS-CoV-2 replication *in vivo* (8, 27–29). Yet, the specific mechanisms by which influenza virus DelVGs disrupt WT replication remain unclear. Most work has focused on the DelVG RNA itself. DelVG RNAs directly interfere with WT virus by competing for access to the viral replication machinery and possibly for packaging into virions (11, 14, 30). DelVG RNAs can also be immunogenic, indirectly interfering with viral replication by triggering innate immune sensing pathways that antagonize WT virus replication (23, 31, 32).

Here, we reveal a completely new actor that interferes with WT virus replication – <u>D</u>elVG-encoded <u>pr</u>oteins (DPRs). We show that DelVGs are translated, producing hundreds of unique proteins. For DelVGs where the deletion remains in-frame, this results in DPRs with native N- and C-termini, but large internal deletions. For DelVGs with out-of-frame deletions, this produces proteins with unique C-termini. DPRs derived from PB2 with native and unique C-termini disrupted replication of WT virus, even demonstrating cross-strain interference. Mechanistic studies showed that PB2 DPRs act as dominant-negative inhibitors that poison the viral replication machinery. Thus, DelVGs have a multipronged inhibitory activity. The RNAs themselves antagonize WT virus, and we now show that the enormous variety of DPRs encoded by DelVGs include proteins that regulate the rate of viral replication.

## Materials and Methods

### Cells, viruses, and plasmids

Experiments were conducted on A549 (ATCC CCL-185), HEK 293T (ATCC CRL-3216), MDCK (ATCC CCCL-34), or MDCK-SIAT-TMPRSS2 cells (a kind gift from J. Bloom (33)). MDCK-SIAT-TMPRSS2 stably expressing PB2 were generated by retroviral gene delivery of a codon-optimized PB2 gene. Cells were maintained in Dulbecco’s modified Eagle’s medium (DMEM; Mediatech 10-013-CV) with 10% FBS and grown at 37° C in 5% CO_2_.

pCDNA3-based vectors for WSN proteins and pBD bi-directional reverse genetics plasmids were previously described (34). pBD-PB2-TRAP-V5 encodes a PB2 gene segment that expresses V5-tagged proteins only when translated from a DelVG (Supplemental File 1). To achieve this, we followed prior work (35) by first duplicating the last 109nt of the PB2 open reading frame and placing it downstream of the native PB2 ORF to create a contiguous packing sequence. We then codon optimized the last 106 nt of the PB2 ORF to avoid direct repeats. Many of the 3’ junctions in DelVGs were located in the last 330 nt of the PB2 ORF. We inserted another copy of this 3’ junction zone downstream of the native PB2 stop codon. This repeat encoded sequence for the V5 epitope tag in all three frames at the 3’ end. It was further modified to remove stop codons upstream of the epitope tag in all three frames. In this way, native PB2 is produced under normal circumstances and can support viral replication, while V5-tagged proteins are only produced if deletions arise that create junctions in the 3’ end of the reporter than maintain an open reading frame with one of the V5 coding sequences. All sequences were confirmed by sequencing.

All virus and virus-derived protein expression constructs are based on A/WSN/1933 except where indicated. Viruses were rescued using the pBD bi-directional reverse genetics system (36). PASTN reporter viruses expressing PA-2A-NanoLuc have been described (37–39). PB2 TRAP V5 virus was generated by replacing wild type PB2 with PB2 TRAP V5. Three different plaque-purified stocks of PB2 TRAP V5 were grown. Virus was amplified on MDCK-SIAT1-TMPRSS2 cells and titered on MDCK cells by plaque assay.

DelVG viruses were created by substituting the PB2 gene segment with a cloned version of a DelVG. PB2 protein was provided in *trans* during transfection of the 293T cells. Virus was amplified and titered using MDCK-SIAT-TMPRSS2 stably expressing PB2.

WSN PB2 TRAP V5 virus and DelVG viruses were confirmed by RT-PCR, sequencing, and western blot.

Transfections were completed using TransIT X2 following the manufacturers’ recommendation.

### Antibodies

V5 proteins were captured using V5 trap agarose beads (Chromotek v5ta). The following antibodies were used for western blot analysis: rabbit anti-PB1 (34), rabbit anti-PB2 (34), rabbit anti-V5 (Chromotek 14440-1-AP), and mouse anti-V5-HRP (Sigma V2260).

### Amplicon-based sequencing of influenza virion RNA using Illumina

A viral stock of WSN or three plaque-purified stocks of PB2 TRAP V5 were treated with 0.25 μg of RNaseA for 30 min at 37°C. Viral RNA was purified with the QIAamp Viral RNA Kit (Qiagen 52904). cDNA was synthesized using the Superscript III Reverse Transcriptase (Invitrogen 18080-044) and primed with MBTUni-12 (5’-ACGCGTGATCAGCRAAAGCAGG-3’). All 8 segments were amplified in a multiplex PCR is MBTUni-12 primer, MBTUni-13 primer (5’-ACGCGTGATCAGTAGAAACAAGG-3’) and Phusion Polymerase (NEB M0530L). PCR products were purified with PureLink PCR Purification Kit (Invitrogen K310002) and eluted in 30µL of Nuclease-free water (Ambion). The RT-PCR procedure was repeated in duplicate for each virus sample. The PCR products were then subjected to next generation sequencing using the Illumina NovaSeq or MiSeq platforms.

### DelVG mapping and quantification in viral stocks

We applied our DelVG-detection pipeline (15) (available at https://github.com/BROOKELAB/Influenza-virus-DI-identification-pipeline). In addition to the standard pipeline, we increased confidence in sequence mapping by using the correlation between replicate sequencing runs of the same sample to calculate the Read Support cutoff (RSC) values (15). DelVG were identified in these filtered dataset using a modified version of ViReMa (15, 40). The presence of repeated sequence in our PB2 TRAP V5 reporter raised the possibility that DelVG assignments may be miscalled. In addition to the standard 25 nt seed used for mapping, we re-ran the analysis using a 52 nt seed, the minimal length needed for unique mapping in the reporter. Only two reads were miscalled and they were excluded from further analysis. The parallel-coordinate maps were generated by adapting an example code (https://observablehq.com/@drimaria/parallel-coordinates-d3-v4) from the D3.js platform.

### Identification of DelVGs in Ribo-Seq Data

Ribosome profiling experiments were previously reported in Tran, Ledwith, et al. 2021 (BioProject PRJNA633047) (41). Data were reanalyzed with a modified version of ViReMa (40) paying attention to read depth at the termini of gene segments and to specifically identify and quantify chimeric ribosome protected fragments (RPFs) spanning the deletion junction of a DelVG.

### Direct RNA sequencing of influenza virions paired mass spectrometry analysis

Long-read sequencing was performed on stocks of WSN that had been generated using a previously-described eight-plasmid reverse genetics system (42), and stocks were used to infect MDCK cells in roller bottles at a multiplicity of infection of 0.01 PFU/cell. After 48 hours, supernatants were clarified by low-speed centrifugation, and influenza virions were concentrated and purified by a two-step ultracentrifugation through a cushion and a gradient of OptiPrep (Sigma) in NTC buffer (100 mM NaCl, 20 mM Tris-HCl pH 7.4, 5 mM CaCl_2_), as previously described (43).

Next, genomic RNA was extracted from purified virions using a Direct-zol RNA Miniprep kit (Zymo) according to the manufacturer’s instructions. 1 µg of freshly purified RNA was prepared for sequencing using a Direct RNA sequencing kit (SQK-RNA002, Oxford Nanopore Technology), with the supplied RT adapter targeting mRNA being replaced with an influenza A virus-specific RT adapter that targets the 12 nucleotides conserved at the 3’ end influenza A virus RNA segments as described previously (44). The modified RT adapter was prepared by annealing primers RTA_FluA (5’-/5PHOS/GGCTTCTTCTTGCTCTTAGGTAGTAGGTTC-3’) and RTA_FluB (5’-GAGGCGAGCGGTCAATTTTCCTAAGAGCAAGAAGAAGCC<u>AGCRAAAGCAGG</u>-3’, replaced sequence underlined). Adapter ligated RNA was sequenced on a MinION nanopore sequencer using FLO-MIN106 flow cell for 48 hours. The resulting fast5 files were basecalled using guppy version 2.3.5.

Nanopore reads were mapped to the WSN genome using minimap2 followed by samtools to create a BAM alignment file (45). Each of the mapped reads within the BAM file were examined for the presence of deletions utilizing the CIGAR alignment string within the BAM file, with the start and end genome coordinates of each deletion extracted. A heatmap was created as a visualization of the frequency of all the deletions greater than 50 nucleotides observed on each genome segment, with the deletion start and end coordinates binned into windows of 50 bases across the genome. The heatmap was created using ggplot (46, 47). Long-read data suggested frequent smaller deletions along the length of each segment of ∼50-100 nt. However, these were not detected at similar frequencies in our amplicon-based sequencing data, and thus it is not clear whether this is reflective of true deletions in the genome or an artifact of the direct RNA sequencing approach. For each deletion observed on each segment, a DelVG segment sequence was created by introducing a corresponding deletion into that sequence from the reverse genetics WSN genome. All of these DelVG segment sequences were then translated in all six frames using EMBOSS-getORF to predict protein sequences with more than 30 amino acids between their start and stop codons (48).

To test whether these predicted proteins could be translated and packaged into virions, we used the six-frame translations of the DelVG sequences to re-query mass spectra previously obtained from purified WSN virions (49). Briefly, this stock of WSN had been grown on MDBK cells and then purified by sucrose gradient ultracentrifugation, either with or without a haemadsorption (HAd) step; its proteins were extracted and digested with trypsin, and liquid chromatography and tandem mass spectrometry (LC-MS/MS) had been performed using a Q Exactive mass spectrometer (Thermo Electron, Hemel Hempstead, UK). The raw data files can be downloaded from MassIVE (http://massive.ucsd.edu/ProteoSAFe/datasets.jsp) using the MassIVE ID MSV000078740 (without HAd: C121116_014.raw, C130214_015.raw, D130107_004.raw; with HAd: C130610_004.raw, C130702_003.raw, C130712_028.raw). Data were analyzed using MaxQuant v.1.6.3.4, using a custom protein database concatenated from several proteomes as described previously (43, 50). The proteomes used were: (i) the *Bos taurus* proteome (UniProt UP000009136, downloaded on the 16^th^ May 2017) and manually edited to remove all repeats of the ubiquitin sequence; (ii) a single repeat of the ubiquitin sequence; (iii) the proteome of our reverse genetics strain of WSN, and (iv) the complete six-frame translations of all of the DelVG sequences identified in the long-read experiment. MaxQuant’s list of common protein contaminants was also included in the search. Protein and peptide false discovery rates were set to a maximum of 1 % using a reverse sequence database. Proteins translated from DelVGs (DPRs) were identified on the basis of peptide spectra that were matched to sequences translated from DelVG segments and which were not found in translations of the reference influenza virus genome segments. The DelVGs identified as encoding DPRs are highlighted on the heatmap by a black circle at the co-ordinates corresponding to the start and end of their genomic deletion.

### Cloning DPRs

RNA was extracted from the supernatant of MDCK cells infected with the PB2 TRAP V5 virus using Trizol (Invitrogen). RNA was reverse-transcribed with Superscript III (Invitrogen) using primers specific to the PB2 segment. cDNA was amplified via PCR using primers annealing to PB2 UTRs that would capture full-length PB2 and DelVGs. Products were cloned into pCDNA3, pBD or the viral RNA expression vector pHH21 and sequenced. To create versions that contained premature stop codons, DPRs cloned into pHH21 were mutated by introducing stop codons in all three frames beginning at the fourth codon of the open reading frame.

### Co-immunoprecipitation

HEK 293T cells were transfected with plasmids expressing NP, PB2-DelVG-V5, PB2, PB1 and PA. Cells were lysed in coIP buffer (50mM Tris pH 7.4, 150 mM NaCl, 0.5% NP40 and 1x protease inhibitors) and clarified by centrifugation. PB2-V5 proteins were captured with V5 trap agarose beads and washed extensively. Precipitating proteins were detected by western blot.

### Polymerase Activity Assay

Polymerase activity assays were performed by transfecting HEK 293T cells with plasmids expressing NP, PB2, PB1 and PA and pHH21-vNA luc expressing a model vRNA encoding firefly luciferase (51). A plasmid expression *Renilla* luciferase was included as a transfection control. Where indicated, PB2-DelVG-V5 proteins were co-expressed to test their inhibitory potential. Cells were lysed 24 hours after transfection in passive lysis buffer (Promega) and firefly and *Renilla* luciferase activity were measured. Results were normalized to the internal *Renilla* control. Protein expression was assessed by western blot.

### Transfection-Infection

HEK 293T cells were seeded into a 6 well plate and transfected the next day with plasmids expressing PB2-DelVG-V5 or an empty pCDNA3 control. Cells were split 1:2 24 h after transfection. 24 h after that, cells were inoculated with the PASTN reporter virus at an MOI of 0.05. Supernatants were harvested 24 h post-infection. Viral titers were measured by infecting MDCK cells and measuring Nanoluciferase activity 8 h later with the Nano-Glo Luciferase Assay System (Promega)(52).

### Co-infection and fitness competition assays

Clonal viruses encoding PB2 DPRs 416/2189 or PB2 195/2223 in place of full-length PB2 were rescued in 293T cells co-expressing WT PB2 protein. PB2 DPR viruses were amplified and titered on MDCK-SIAT-TMPRSS2 stably expressing PB2. Co-infections were performed by inoculating MDCK cells with WT PASTN virus (MOI = 0.05), and where indicated PB2 DPR viruses were also added (MOI = 0.05 or 0.1). Supernatant was recovered and yield of WT virus was measured via Nano-Glo viral titer assay (37).

*HA* segments expressing PB2 DPRs were designed as before (53) and cloned into pHW2000. All upstream start codons present in the packaging signals were mutated and sequence for PB2 416/2189 or a mutant containing a premature stop codon was inserted. Viruses encoding the artificial *HA* segment were rescued, amplified and titered on cells expressing HA protein. Competition experiments were performed by infecting HA-expressing MDCK cells at an MOI = 0.01 with a 1:1 mixture of viruses with WT PB2 416/2189 or the premature stop mutant. Virus was harvested 72 hpi. RNA was extracted from the inoculum mix and output virus and was quantified with genotype-specific primer pairs by RT-qPCR. WT was detected with 5’- CTCGCACTCGCGAGATACTC-3’ and 5’-CCAAAAGCACCAGGATCATTG-3’ while the premature stop mutant was detected with 5’-CTCGCACTCGCGAGATATAG-3’ and 5’- AGACCAAAAGCACCAGGATCCCTA-3’. Plasmids bearing each variant were used to generate a qPCR standard curve, against which a linear regression was used to back-calculate normalized copy number values for each molecular species and correct for relative inefficiencies in qPCR. Inoculums were generated from two independent virus rescue experiment. Each inoculum was then used for two independent competition experiments. Each qPCR value was calculated as the mean of technical triplicates. A competitive index was calculated for each of the four trials by dividing the mutant:WT ratio following infection by the mutant:WT in the inoculum.

### Statistics

Experiments were performed in at least biological triplicate, with each including at least three technical replicates. Results are presented as grand mean of three biological replicates ± standard error, or mean of a representative biological replicate ± standard deviation, as indicated. Statistical significance of pairwise comparisons were made with a two-tailed Student’s t-test or one-sample t-test. Multiple comparisons were assessed by ANOVA with *post hoc* Dunnett’s multiple comparisons test. The correlation between input RNA and ribosome-protected fragments was determined using a Spearman’s correlation coefficient. χ^2^ tests were used to test for deviation from an expected distribution.

## Results

### Translation of DelVG mRNAs

DelVGs are replicated and transcribed, but the degree to which they are translated is unknown. We performed amplicon-based sequencing of virion-associated RNA from A/WSN/33 (WSN; H1N1) stocks to identify and characterize the DelVGs present in our viral stocks. As DelVGs contain large internal deletions, our analysis pipeline explicitly allowed discontinuous mapping of reads for precise identification of deletion junctions (15). In addition to full-length vRNA, DelVGs originating from all genomic segments except *HA* were identified (Fig 1a and Fig S1a). Of the 137 DelVGs detected, 86% were derived from the polymerase gene segments (Supplemental Table 1), consistent with prior work showing that these larger segments are more likely to be the source of DelVGs (13, 16–21). DelVG breakpoints were heavily concentrated near the termini, with ∼90% starting within the first 400 nt of a fragment and rejoining with the last 450 nt (Fig 1a-b, S1a-b, Supplemental Table 1). While we represent DelVGs in cRNA orientation, it is not known whether deletions were formed during cRNA or vRNA synthesis. Further, the DelVGs that were detected are not necessarily reflective of all of those that were made, but rather those that were maintained in the viral population over multiple rounds of replication (14).

**Figure 1.**
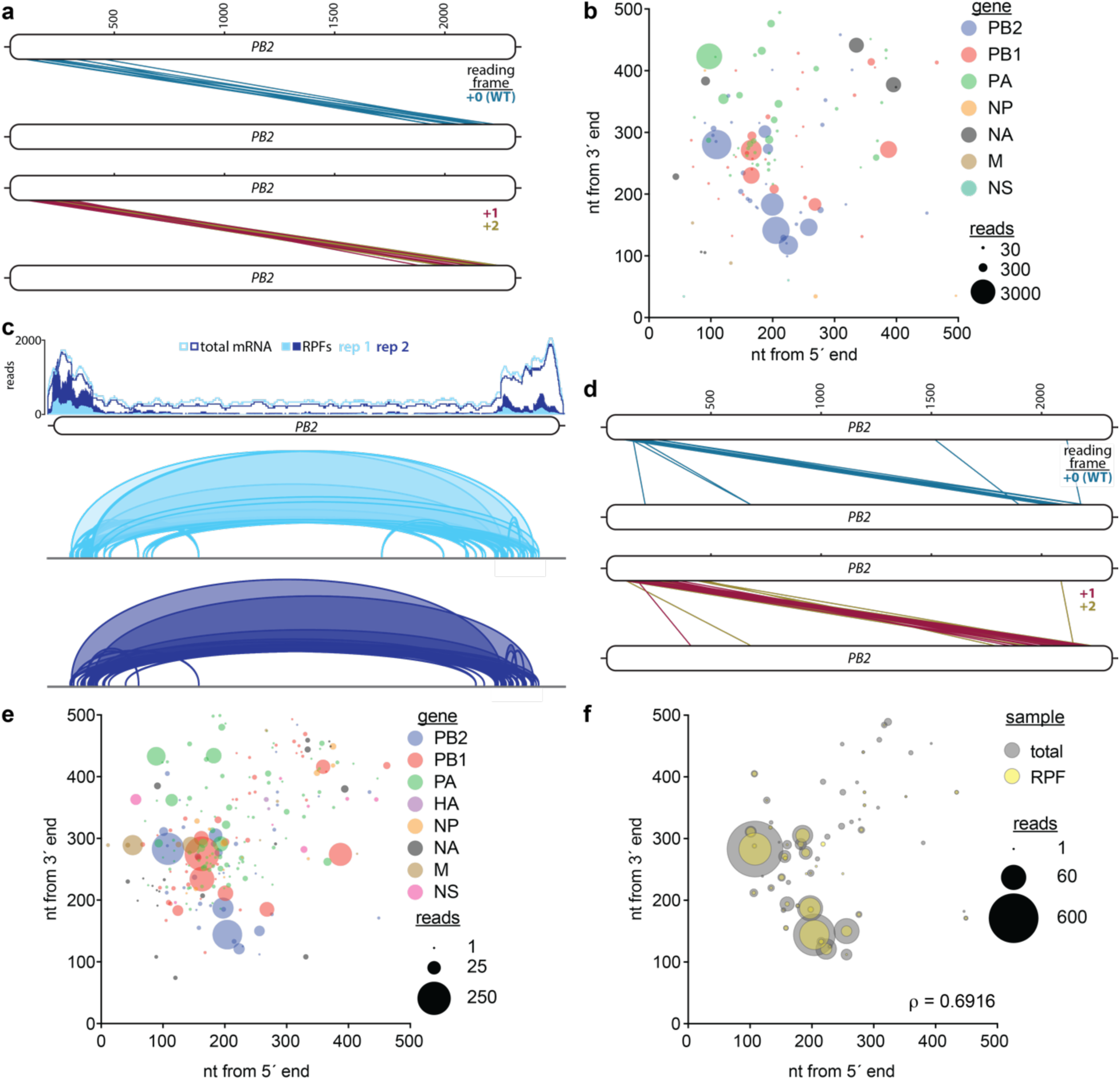
Transcription and translation of deletion-containing viral genomes (DelVGs). **(a)** Virions contain DelVGs with protein-coding potential. DelVG were identified by amplicon-based sequencing of genomic RNA purified from WSN virions and visualized by parallel coordinate mapping. *PB2* is shown as plus-sense cRNA, and each *PB2* DelVGs is depicted by a line connecting the 5’ (top) and 3’ (bottom) ends of each deletion. The reading frame downstream of the deletion was predicted, revealing the potential to produce internally deleted PB2 variants in the +0 frame or chimeric proteins in +1 and +2 frames. (**b)** Junctions and read depth were determined for all DelVGs and plotted as a function of their positions from the ends of the respective viral gene. Note that none of the DelVGs in this sample mapped to *HA*. **(c)** DelVGs are translated. Ribo-Seq data obtained from cells infected with the same viral stocks as above were reanalyzed to allow for discontinuous mapping of ribosome-protected fragments (RPFs). *top,* Read depth of total mRNAs (lines) and RPFs (filled) were mapped to *PB2* in two independent replicates. *bottom,* RPFs containing deletion junctions are shown as arcs, where line width reflects read depth. (**d)** Parallel coordinate mapping of PB2 RPFs containing deletion junctions, performed as in (a). (**e)** Junctions and read depth were determined for all RPFs containing deletions and plotted as a function of their positions from the ends of the respective viral gene. (**f)** The abundance of deletion-containing fragments from *PB2* is correlated between total RNA and RPFs. Data are composite of two independent Ribo-Seq experiments. Spearman’s correlation coefficient (r) between total and RPF read depth for each DelVG is shown. See also Figure S1-S2 and Supplemental Tables 1-2 for remaining gene segments, bubble plots showing the entire gene length, and predicted protein products.

Analysis of DelVG sequence revealed that 30% retained the original reading frame after the deletion (Fig 1a; Supplemental Table 1), raising the possibility that mRNAs transcribed from DelVGs have the potential to produce protein products that are fusions between the N- and C-termini of the parental protein. The remaining DelVGs shifted into the +1 or +2 reading frame downstream of the junction. Transcripts from these would code for proteins with a native N-terminus followed by entirely new protein sequence downstream of the junction until encountering a stop codon that is normally out of frame. For PB2 DelVGs in our stock, we predicted this would result in translation of up to 17 amino acids from an alternative reading frame downstream of the native N-terminus (Supplemental Table 1).

This coding potential suggests that DelVGs may be translated. To determine if DelVGs mRNAs are translated during infection, we re-analyzed ribosome profiling (Ribo-Seq) data we had previously obtained from lung cells infected with WSN (41). High-throughput sequencing of total RNA from infected cells showed high read coverage at the termini of *PB2,* consistent with the presence of DelVGs in our viral stocks (Fig 1c). This pattern was also seen in ribosome-protected fragments. Mapping of ribosome-protected fragments on *PB2* identified 50 unique DelVG junctions (Fig 1c, bottom), which could only occur if DelVGs were actively undergoing translation. Over 300 unique DelVGs derived from all 8 segments were detected in ribosome-protected fragments across two biological replicates (Fig S2a, Supplemental Table 2). A similar distribution of DelVGs was present in the total RNA extracted from these cells (Fig S2b). The reading frame downstream of the deletion junctions in ribosome-protected fragments did not differ significantly from an equal distribution across all three frames (χ^2^ *p*>0.05, Supplemental Table 2). Most of the predicted DelVG-encoded proteins were less than 250 amino acids in length (Fig S2c). Reading frame usage showed no obvious correlation with deletion size or the position of deletion junctions for both ribosome-protected fragments and total RNA (Fig 1d, S2b). As in our viral stocks, deletion junctions were enriched near the termini of gene segments (Fig 1e, S2d-e). The abundance of specific PB2 DelVGs in total RNA versus ribosome-protected fragments is positively correlated (Fig 1f). This correlation may be underestimated by our data, as short-read sequencing of total mRNA has inherent length-dependent bias that over-represents smaller fragments, which is absent from the Ribo-Seq approach. This suggests that the probability of translation is a function of concentration with no obvious preference for a particular type of DelVG mRNA.

DelVGs arise naturally during influenza virus infections in humans and animals (18–21, 23). Analysis of sequencing results from human influenza virus infections revealed DelVGs with protein-coding potential in challenge studies with A/Wisconsin/67/2005 (H3N2) and in naturally acquired infections with A/Anhui/1/2013 (H7N9) (19, 21). DelVGs from patients were primarily derived from the polymerase segments. Human-derived PB2 DelVGs from both studies encoded potential protein products in all three frames, and several of these were of similar sequence to those detected in our viral stocks (Fig S3 and Supplemental Table 3). Thus, DelVGs with protein coding potential arise *in vitro* and *in vivo* and their mRNAs are translated into proteins, some of which contain entirely new C-terminal sequences derived from alternative reading frames.

### Cryptic viral proteins produced by DelVG mRNAs

Influenza A virus encodes at least 14 canonical proteins (54). To test for the presence of non-canonical DelVG-encoded proteins, we combined direct RNA sequencing of viral genomes with mass spectrometry proteomics. Direct RNA sequencing covers entire genome segments in a single continuous read without copying into cDNA or amplification allowing for direct detection of deletions. Direct RNA sequencing of virion-derived genomes identified DelVGs with large internal deletions in each genome segment that retained the terminal ∼200 nt (Fig 2a), mirroring our short-read data. These DelVG sequences, along with the full-length viral genome, were then subjected to a complete 6-frame translation *in silico* to create a search database of hypothetical proteins that could be translated from a wide variety of different DelVGs. Mass spectrometry analysis was performed on proteins extracted from purified virions. Mapping spectral matches to the 6-frame database identified peptides derived from translation of both the full length reference mRNAs and of DelVG mRNAs (Fig 2a-b and Supplemental Table 4). A large constellation of DelVG-encoded proteins (DPRs) were detected, derived from 7 of the 8 genome segments, ranging from a big DPR of 526 amino acids encoded by a *HA* DelVG to a little DPR of only 18 amino acids that shared an N-terminus with PB1-F2 (Fig 2b). Some of these peptides appear to have originated from proteins using alternative initiation codons (Supplemental Table 4). Within virions, DPRs translated from DelVGs of *M* produced the highest number of unique peptides. Whether this distribution reflects overall DPR abundance, stability, or the fact that we analyzed proteins packaged into virions remains to be determined. Together, our results show that not only do DelVGs have coding potential, but they are also translated and produce unique proteins, some of which are incorporated into virions.

**Figure 2.**
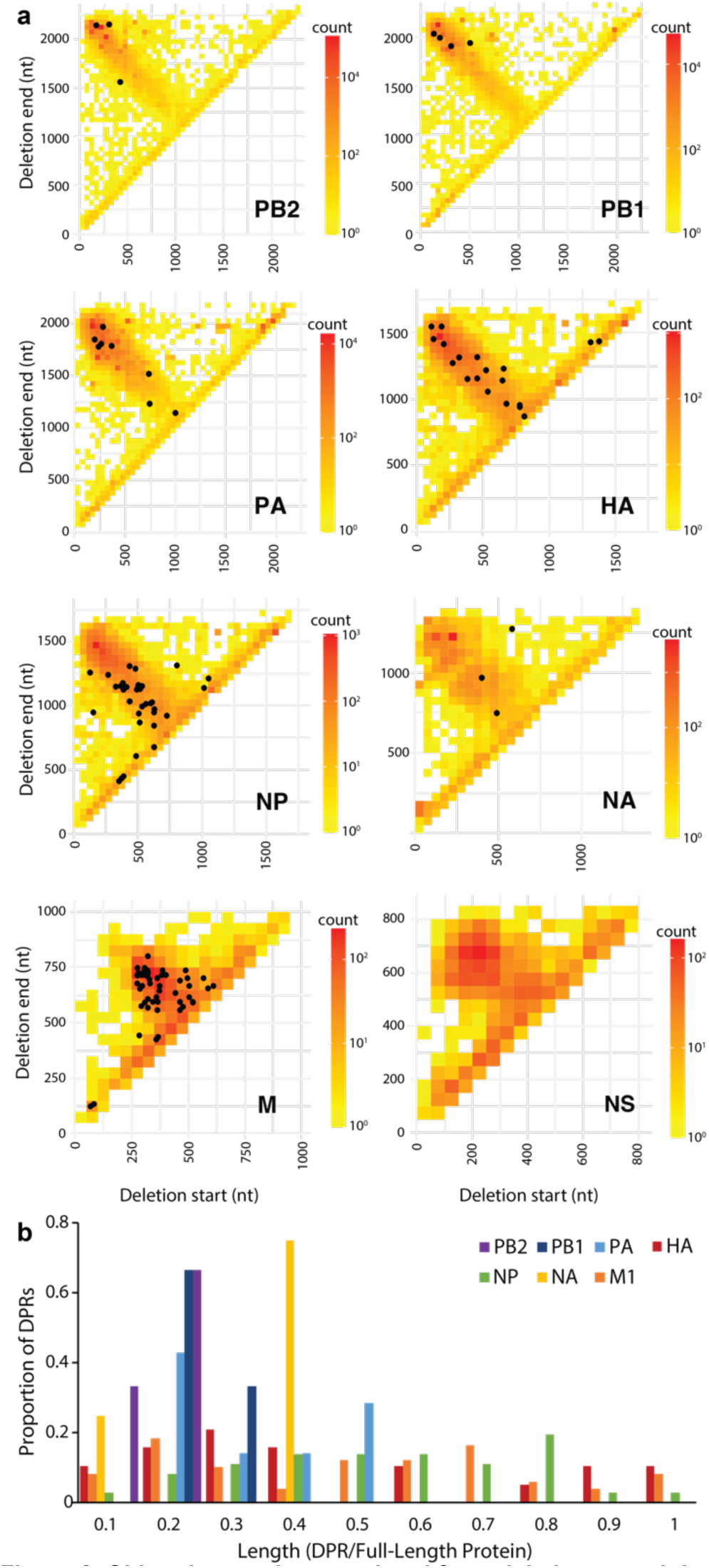
Chimeric proteins translated from deletion-containing viral mRNAs. **(a)** DelVGs were identified by direct RNA sequencing of virion-derived viral genomes. Deletion junctions for each of the 8 genomic segments were binned into 50 nt groups and plotted on heatmaps to represent their abundance. These DelVGs were used to create a query database for detection of DelVG-encoded proteins. Proteins extracted from purified virions were subject to mass spectrometry. Identified chimeric peptides are indicated by black dots mapped onto to the DelVGs that encoded them. (**b)** Histogram showing the lengths of DPRs encoded by DelVGs detected by direct RNA sequencing compared to the length of the corresponding protein encoded by a full-length segment.

### DelVG-encoded proteins arise *de novo*

Our viral stocks were produced via plasmid-based reverse genetics (55), and in theory should have started as a clonal population containing full-length genomes. The DelVGs and DPRs that we identified presumably arose as our viruses replicated to produce working stocks. To elucidate the emergence and translation of DelVGs and the potential function of these proteins, we designed the influenza virus reporter PB2 V5 TRAP (Fig 3a). The majority of *PB2* DelVGs contain 3’ junctions within the last 300 nt of the cRNA (Figs 1, 2) and so we created a virus that would identify deletions ending in this region. To do this, we inserted an additional copy of this 3’ junction zone downstream of the native *PB2* stop codon. The repeated region was modified to remove stop codons and to encode V5 epitope tags in all three reading frames. In this way, the PB2 V5 TRAP virus produces native PB2 from its full-length genomes, while V5-tagged DPRs would be produced by DelVGs with a break-point placed in any frame of the repeated junction zone.

**Figure 3.**
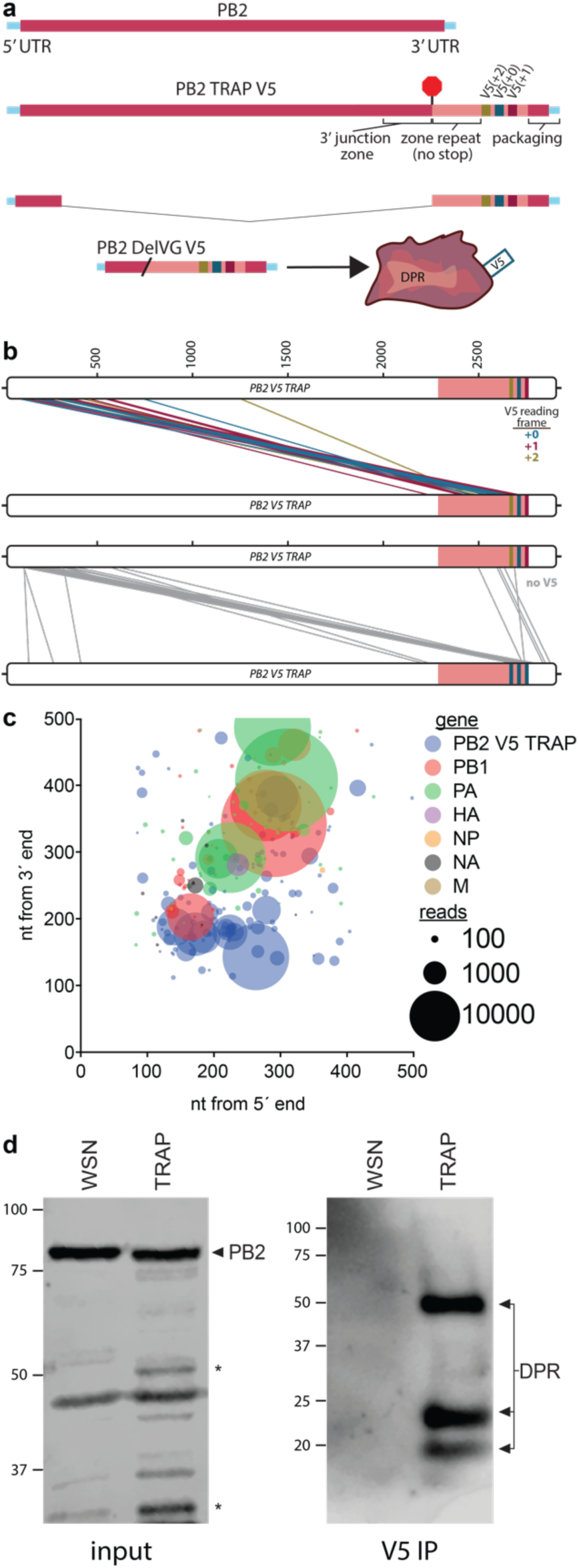
DelVG-encoded proteins are expressed during infection. **(a)** Diagram of the *PB2* gene in a A/WSN/33 reporter virus designed to capture expression of DelVG-encoded proteins. Sequence from the 3’ junction zone of DelVGs was repeated downstream of the native stop codon. Stop codons were purged in all three frames from the repeated sequence to enable expression of a V5 epitope tag on proteins translated from this region. (**b)** Parallel coordinate mapping of DelVGs identified by amplicon-based sequencing the PB2 TRAP V5 stock #1. DelVGs with the potential to express V5-tagged proteins (*top*) and the remaining DelVGs (*bottom*) are shown. (**c)** Junctions and read depth were determined for all DelVGs in three independent plaque-purified stocks and plotted as a function of their positions from the ends of the respective viral gene. Note that none of the DelVGs in this sample mapped to *NS*. **(d)** Expression of DelVG- encoded proteins (DPRs) in infected cells. Cells were infected with WT or PB2 V5 TRAP virus. Infected-cell lysate was probed with anti-PB2 antibody by western blot, while epitope-tagged DPRs were immunoprecipitated and detected with anti-V5 antibody. For (a) and (b) See also Figure S4, S4 and Supplemental Table 4.

Recombinant PB2 V5 TRAP virus was rescued and used to generate plaque-purified stocks to ensure we began with a clonal population. Importantly, only viruses encoding a full-length PB2 gene should be able to replicate during plaque purification. Three clones of reporter virus were selected, amplified as viral stocks, and DelVGs were identified by amplicon-based sequencing of vRNA. Over 350 distinct DelVGs were detected in all segments except *NS* (Fig S4a, Supplemental Table 3). Again, DelVGs were enriched in the polymerase genes. Patterns of DelVG junctions paralleled those present in WT WSN stocks (Fig 3b-c, S4a-b), except that in the PB2 V5 TRAP reporter 3’ junctions were relocated downstream to the repeated junction zone as anticipated (Fig 3b). DelVGs encoding >100 potential V5-tagged proteins were identified (Supplemental Table 5). The +1 reading frame was overrepresented (χ^2^ < 0.05), but whether this bias represents a preference in PB2 DPR production or is a consequence of the reporter design is not clear.

Cells were infected with WT or the PB2 V5 TRAP virus. Lysates were probed for PB2, revealing multiple lower molecular weight proteins (Fig 3d). To confirm that these were DelVG-encoded, antibodies that recognize the V5 tag were used to immunopurify proteins from the lysates and to detect the recovered proteins by western blot. Three prominent bands were detected exclusively in cells infected with the reporter virus, including a 50 kDa protein that was also clearly detected in the total lysate. These data from plaque-purified virus demonstrate that DPRs are produced from DelVGs that arise *de novo* during viral growth. Our sequence analysis, proteomics and experimental validation provide multiple lines of evidence that reveal DPRs as a new class of viral proteins.

### PB2 DPRs are dominant negative inhibitors of polymerase activity and viral replication

The function of DPRs, if any, was not known. To investigate this, we cloned PB2 DPRs from our reporter virus for further study. Of nine randomly cloned PB2 DelVGs, eight had protein coding potential (Fig 4a, Supplemental Table 6). The ninth, PB2 32/2143, had a premature stop after the first 4 codons; this then served as a negative control in functional assays allowing us to isolate the impact of DelVG RNA from that of DPRs. Five of these potential proteins from PB2 V5 TRAP have equivalent versions when mapped back to WT *PB2 (*Fig 4a). As PB2 is an essential subunit of the viral polymerase, we assessed the impact of PB2 DPRs on polymerase function. PB2 DPRs were co-expressed in polymerase activity assays. Polymerase activity was significantly reduced by all DPRs, except the negative control PB2 32/2143 (Fig 4b). Thus, proteins encoded by PB2 DelVGs interfere with polymerase activity.

**Figure 4.**
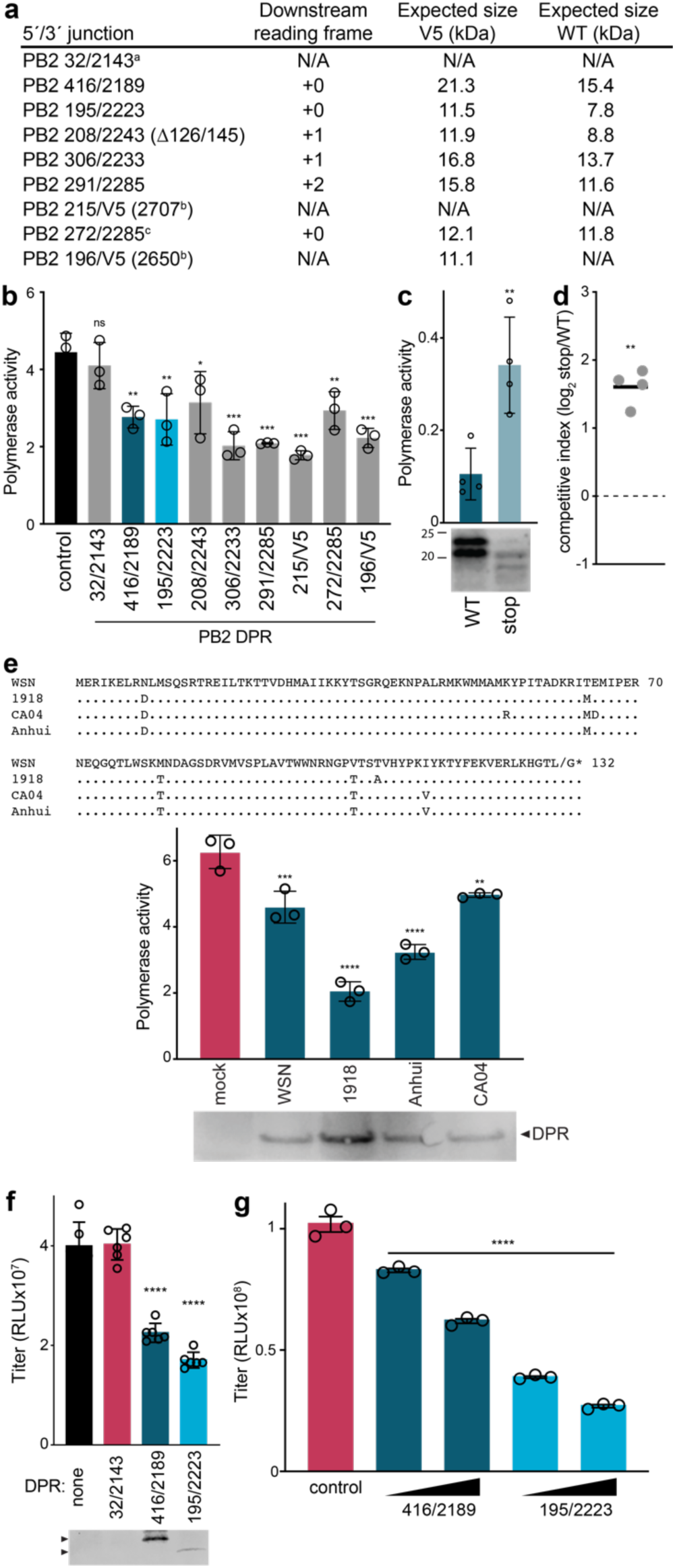
DPRs are dominant-negative inhibitors of the influenza virus polymerase and viral replication. **(a)** DelVGs cloned from cells infected with the PB2 TRAP V5 reporter virus. Deletion junctions were mapped back to WT PB2 and used to name each DelVG. Numbering is based on full-length *PB2* cRNA and indicates the last nucleotide of the upstream portion and the first nucleotide of the downstream sequence. The expected size of the V5-tagged DPR from the reporter virus is indicated as is the predicted size if this deletion arose in a wild-type virus. PB2 208/2580 (Δ126/145) contains an additional deletion. ^a^Degenerate sequence at the junction precludes precise mapping, with the junction spanning 25-32/2136-2143. ^b^The 3’ junction is within the V5 epitope coding sequence and does not have analogous position in WT virus. ^c^The equivalent DelVG in a WT background lacks a stop codon, making it unclear if it would produce a protein. **(b)** Polymerase activity assays were performed in 293T cells in the presence or absence of the indicated DPRs. PB1, PB2, PA, NP and DPRs are all derived from WSN. Data are normalized to co-transfected *Renilla* luciferase. (**c)** Polymerase activity assays were performed as in (b) in the presence of PB2 416/2189 or a version containing premature stop codons. DPR expression was detected by western blot. (**d)** Fitness competition assays were performed by coinfecting with a virus encoding PB2 416/2189 and a version containing a premature stop codon. Genotypes were quantified by RT-qPCR and a competitive index was calculated for the premature stop mutant versus WT following coinfection. **(e)** Polymerase interference by heterotypic DPRs. Polymerase activity assays in the presence or absence of the PB2 416/2189 DPR were performed as in (b), expect the DPR was cloned from the indicated viral strain. DPR sequence is shown at the top and expression was confirmed by western blot. (**f-g)** DPRs suppress viral replication. (**f)** The indicated DPRs were ectopically expressed in cells prior to infection with an influenza reporter virus. Supernatants were harvested 24 hpi and viral titers were measured. DPR expression was assessed by western blot. (**g)** Cells were co-infected with a wild-type reporter virus and a clonal virus encoding the indicated DPR in place of *PB2*. Replication of the wild-type virus was measured by titering the supernatant via a luciferase-based assay. For b-f data are mean of a representative biological replicate of n=3-6. For g, data are shown as the grand mean of n=3 independent biological replicates, each performed in technical triplicate. Statistical significance was assessed by Student’s t-test (c), a one-sample t-test (d) or for others, using an ANOVA with *post hoc* Dunnett’s multiple comparisons test. * < 0.05, ** < 0.01, *** ≤ 0.001. **** < 0.0001.

We tested the cumulative effects of both the DelVG RNA and DPR on polymerase activity and viral replication. We focused on PB2 416/2189 as it was the most abundant DPR amongst those that we cloned from our reporter stock. Similar DelVGs were also identified in our WSN stock and Ribo-Seq data (Supplemental Table 1, 2, 4). Expressing the PB2 416/2189 vRNA dramatically reduced polymerase activity compared to the control (Fig 4c). This inhibitory activity was partially dependent on the encoded DPR, as polymerase activity was partially restored in the presence of a mutant vRNA containing premature stop codons that reduce DPR expression. Curiously, despite three stop codons in different frames at the very beginning of the open reading frame, this mutant still expressed low amounts of proteins, possibly by reinitiating at the methionine at position 11 (Fig 4c). We further tested the contribution of DelVG RNA versus DPR in the context of viral replication. We engineered viruses where the *HA* gene was replaced with PB2 416/2189, or by a mutant PB2 416/2189 with a stop codon at position 20. These viruses were grown on cells that supplied HA protein *in trans*. The relative fitness of each genotype was measured by co-infecting cells with an equal mixture of DPR-expressing and premature-stop mutant viruses. A competitive index was calculated by comparing genotype ratios after competition to those in the inoculum. The mutant virus with the truncated DPR outcompetes virus expressing WT PB2 416/2189 (Fig 4d), indicating that DPR expression reduces fitness. While DelVGs were known to interfere with replication (6), these data now show that the translated DPRs themselves also directly contribute to this inhibition.

The sequence for WT PB2 416/2189 is highly conserved amongst influenza A viruses, raising the possibility that a wide variety of strains, including those in naturally-occurring infections, could produce DPRs with inhibitory activity (Fig 4e). To test this, polymerase activity assays were performed with the WSN-derived polymerase and NP in combination with PB2 416/2189 DPRs cloned from different primary isolates of influenza A virus. We detected heterotypic inhibition of WSN polymerase using DPRs derived from the 1918 H1N1 pandemic virus, the 2009 H1N1 pandemic virus, and an emerging H7N9 virus (Fig 4e). The sequence and deletions in several of the potential DPRs identified in human H3N2 and H7N9 samples were similar to the inhibitory DPRs we tested here (Supplemental Table 3), suggesting that inhibitory activity of DPRs may be shared across influenza virus strains.

Given the ability of DPRs to disrupt polymerase activity, we tested their impact on viral replication. DPRs were expressed in cells prior to infection. Pre-expressing PB2 416/2189 or PB2 195/2223 reduced virus titers by 50% (Fig 4f). In contrast, the PB2 32/2143 DelVG that contains a premature stop codon had no effect, yielding viral titers indistinguishable from controls. To exclude the possibility that the pre-expression process artifactually altered replication, we utilized influenza virus itself to deliver the DPR. We created viruses encoding PB2 416/2189 or PB2 195/2223 in place of full-length PB2. These viruses were propagated and titered on cells constitutively expressing native PB2 protein. DPR viruses were used for co-infection experiments with a WT WSN NanoLuc reporter virus (37). Titering samples using the NanoLuc reporter allowed us to uniquely measure replication of the WT virus, and not the DPR virus (52). Co-infection with increasing amounts of the DPR virus showed a dose-dependent inhibition of WT WSN, reducing titers by up to 75% (Fig 4g). Our data show that DPRs inhibit replication and that viruses encoding DPRs function as classic defective interfering particles.

### PB2 DPRs are dominant negative inhibitors of polymerase assembly

The influenza virus polymerase assembles into a functional trimer containing PB1, PB2 and PA. The N-terminal 37 amino acids of PB2 form three α-helices that intertwine with the C-terminus to PB1 (56). Most of our PB2 DPRs retained this portion of the full-length protein. This region is sufficient for PB1 binding (56, 57), raising the possibility that PB2 DPRs might interact with PB1. To test this, cells were infected with the PB2 V5 TRAP virus and DPRs were immunopurified from cell lysates. Blotting revealed co-precipitation of PB1 when V5-tagged DPRs were present, but not during infection with WT virus (Fig 5a). Interactions between PB1 and PB2 DPRs were investigated further by expressing only these two proteins in cells. Again, PB1 co-precipitated with the PB2 DPRs 416/2189 and 195/2223 (Fig 5b). Thus, not only do these proteins interact, PB2 DPRs are likely directly engaging PB1 without the need for other viral proteins.

**Figure 5.**
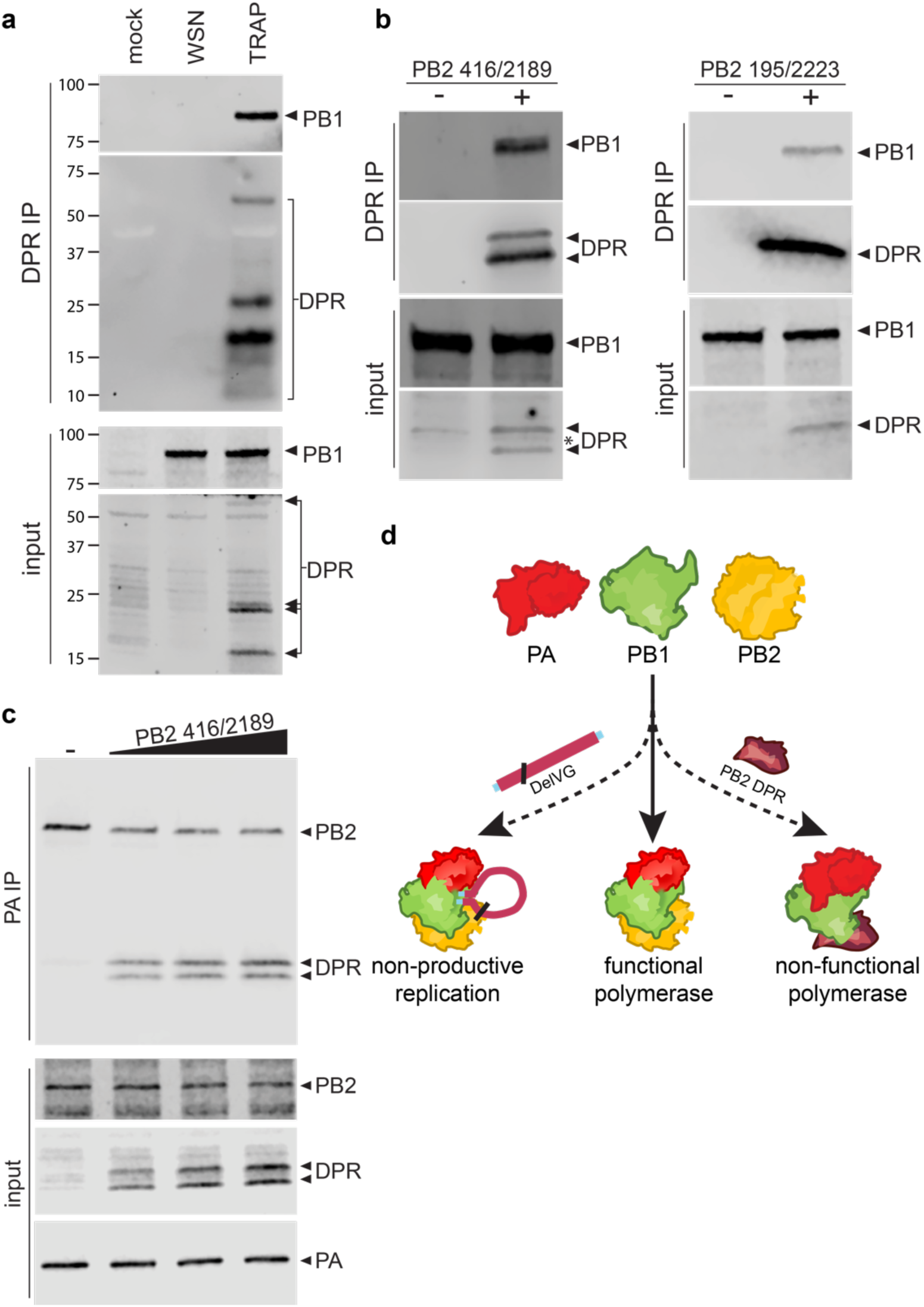
DPRs compete for assembly of function viral polymerase. **(a-b)** PB2 DPRs interact with PB1. (**a)** Lysate were prepared from cells infected with the PB2 TRAP V5 virus and DPRs were immunopurified via their V5 tag. DPRs and co-precipitating proteins were detected by western blot. (**b)** The viral polymerase proteins and NP were expressed in the presence or absence of the indicated DPR. DPRs were immunopurified from cell lysates and proteins were detected by western blot. (**c)** PB2 DPR competes for polymerase assembly. The viral polymerase proteins and NP were expressed in the presence of increasing amounts of the PB2 DPR. Polymerase assembly was measured by immunoprecipitating PA and probing for PB2. (**d)** A dual mechanism for the inhibitory activity for DelVGs and the DPRs they encode.

Interfering with trimer assembly disrupts polymerase activity (56–59). Given that PB2 DPRs bind PB1 and impair polymerase function, a potential mechanism is that they competitively inhibit trimer formation. We tested this with an RNP assembly assay where the polymerase proteins and NP are expressed in cells, immunopurified via PA, and blotted for PB2. PA does not form a stable, direct interaction with PB2. Therefore, any co-precipitated PB2 must be part of a trimeric complex and reflect successful polymerase assembly. PB2 was readily co-precipitated with PA under control conditions (Fig 5c). However, expressing increasing amounts of PB2 416/2189 displaced WT PB2, and instead the DPR was co-precipitated (Fig 5c). These data demonstrate that PB2 DPRs are dominant-negative inhibitors of trimeric polymerase assembly, establishing the mechanism by which they reduce in polymerase activity and viral replication.

## Discussion

DelVGs are aberrant replication products with the potential to interfere with the replication of WT virus. DelVGs contain large internal deletions while retaining terminal sequences necessary for replication, transcription and packaging by viral machinery. We now reveal that influenza virus DelVGs template synthesis of hundreds of cryptic viral proteins. These DPRs usually contain native N-termini fused to C-termini derived from all three reading frames, creating novel protein products. Functional assays demonstrate that DPRs make key contributions to the inhibitory potential of DelVGs and their interference with replication of influenza virus. Given that DelVGs are produced by most RNA viruses, our results suggest DPRs have the potential to dramatically expand the functional space of proteins encoded by a diverse array of human pathogens.

PB2 DPRs disrupt virus replication by competing with WT PB2 for binding to PB1 (Fig 5). Only the N-terminal 37 amino acids of PB2 are required to form a PB1-binding domain and disrupt polymerase function (56, 57). We identified ∼270 distinct DelVGs across our samples that code for this domain. Thus, any PB2 DPR that contains this region and is stably expressed has the potential to function as a dominant-negative inhibitor. The inhibitory potential of PB2 DPRs may be further aided by the fact that DelVGs are often present at much higher levels than their WT counterparts. We identified analogous DPRs from PB1. The N-terminal ∼12 amino acids of PB1 encompass the minimal binding domain for PA (60), and were found in every PB1 DelVG that we identified. This region expressed as a fusion protein, or even delivered as a peptide, blocks binding of full-length PB1 to PA (58, 61). PB1 DPRs that retain the native C-terminus may also disrupt trimer assembly, as they would contain the PB2 binding domain (62). These suggest that PB1 DPRs would also inhibit replication by disrupting polymerase assembly. Many of the DPRs from other gene segments retained defined functional domains. It will be important to determine if they impact replication as well.

DPRs join a growing collection of non-canonical protein products made during influenza virus infection. The virus exploits splicing, leaky scanning and frame-shifting to encode multiple protein products (63). Recent work has also revealed additional, perhaps more fortuitous mechanisms to produce proteins. Start-snatching involves translation initiation in the host-derived regions of viral mRNAs, upstream of the normal start codon. The resultant products create N-terminal extensions to native viral proteins or novel polypeptides when initiation is out of frame, and some of these affect virulence *in vivo* (64). Leaky scanning expresses cryptic viral epitopes from the *M* and *NS* segments that drive potent CD8 T cell response during infection (65, 66). Translation products have also been detected from a conserved open reading frame of the negative strand of genome segments (67). While these negative strand-encoded products are detected by T cells as part of immunosurveillance, their impact on viral replication is currently an open question. Our large-scale detection of DPRs here complements earlier reports that identified specific proteins encoded by individual DelVGs. *In vitro* translation of purified DelVG mRNAs identified peptides from PB2 and PA, but no function was assigned (68, 69). More recently, a 10 kDa protein from a *PB2* DelVG was identified and shown to activate innate immune signaling (70). Our data indicate DPRs are common components of influenza virus infection. It should be noted that our ability to detect DPRs is dependent upon the DPR being sufficiently tolerated by the virus to be maintained in the population. Exceptionally potent DPRs would suppress WT virus to such an extent that they would be lost. Thus, our identification likely under-samples the total sum of DPRs that are produced and the full spectrum of their inhibitory potential. We have defined a discrete function for some of the PB2 DPRs in disrupting polymerase assembly. It is tempting to speculate that they may impact replication not just at the cellular level, but also at the population level by establishing a negative feedback loop that tempers the outgrowth of inhibitory DelVGs. The discovery of DPRs opens an entire new collection of protein chimeras with the potential to alter population fitness, pathogenicity or present novel immunogenic epitopes.

DelVGs are produced by a diverse range of RNA viruses and have been shown to impact pathogenesis during human infections with influenza virus, respiratory syncytial virus, and SARS-CoV-2 (9, 24, 71, 72). This raises the possibility that DPRs may be a generalizable feature of RNA virus replication. By extension, exploiting DelVGs and the specific DPRs they encode may offer a universal therapeutic strategy for these infection. In summary, we have shown that DelVGs interfere with viral replication via a one-two punch – both the RNA itself and the encoded DPR combine to antagonize WT virus.

## Data Availability

Raw data underlying results presented in the figures can be found in Supplemental Tables 7 and uncropped blots are present in Fig S5. Accession numbers for all deposited data are in Supplemental Table 8. Ribo-Seq data are available in BioProject PRJNA633047. Sequencing data from viral stocks are accessible as BioProject PRJNA957839. Direct-RNA sequence data are in BioProject PRJNA1051370. Mass spectrometry spectra are accessible as MassIVE ID MSV000078740.

## Acknowledgements

We thanks members of the Mehle lab for comments on the manuscript and Daniel Mair for technical assistance with sequencing.

