## Supplemental Figure 1 for "Cryptic proteins translated from deletion-containing viral genomes dramatically expand the influenza virus proteome"

**Figure S1. Deletion-containing viral genomes (DelVGs) from viral stock. Associated with Figure 1a-b.**
**(a)** DelVG were identified by sequencing genomic RNA purified from WSN virions and visualized by parallel coordinate mapping. Genes are shown as plus-sense cRNA, and each DelVGs is depicted by a line connecting the 5´ (top) and 3´ (bottom) ends of each deletion. (**b)** Junctions and read depth were determined for all DelVGs and plotted as a function of their positions from the ends of the respective viral gene. These are the same data as in Fig 1b, but scaled to show the full length of the gene segment. Note that none of the DelVGs in this sample mapped to *HA*. See also Supplemental Table 1 for underlying data.

### Fig S1

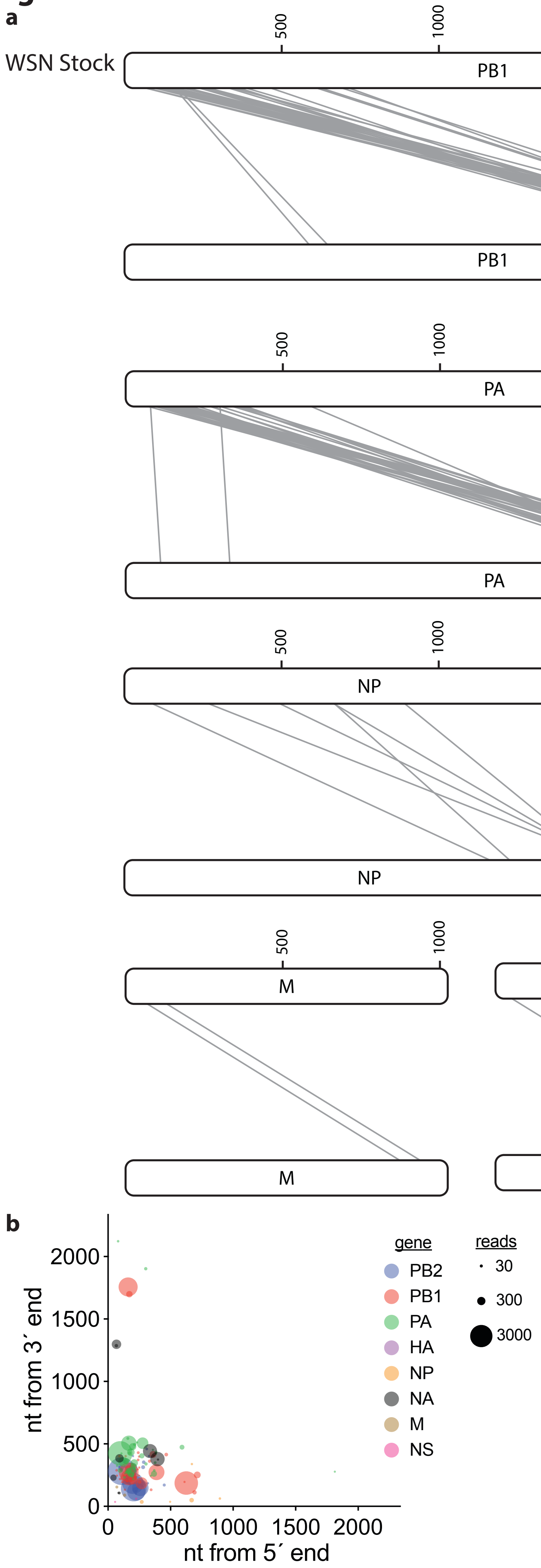
a
WSN Stock
PB1
PB1
PA
PA
NP
NP
NA
NA
M
M
NS
NS
500
1000
1500
2000
b
gene
PB2
PB1
PA
HA
NP
NA
M
NS
reads
30
300
3000
nt from 3´ end
nt from 5´ end
0
500
1000
1500
2000
