## Supplemental Figure 2 for "Cryptic proteins translated from deletion-containing viral genomes dramatically expand the influenza virus proteome"

**Figure S2. Ribo-Seq data reveal transcription and translation of deletion-containing viral genomes (DelVGs). Associated with Figure 1c-f.**

**(a)** Ribo-Seq data were reanalyzed to allow for discontinuous mapping of ribosome-protected fragments (RPFs). Genes are shown as plus-sense cRNA, and each DelVGs is depicted by a line connecting the 5´ (top) and 3´ (bottom) ends of each deletion. (**b)** DelVGs were identified in the total RNA input controls from Ribo-Seq and mapped as in (a). *PB2* was further separated based on predicted reading frames downstream of the deletion. (**c)** Predicted protein length for DelVGs detected in total RNA and RPFs. (**d)** Junctions and read depth were determined for all RPFs containing a deletion and plotted as a function of their positions from the ends of the respective viral gene. These are the same data as in Fig 1e, but scaled to show the full length of the gene segment. **(e)** Junctions and read depth were determined for all DelVG in total RNA input controls and plotted as a function of their positions from the ends of the respective viral gene. For all panels, data are composite of two independent Ribo-Seq experiments. See also Supplemental Table 2 for underlying data.

### Fig S2

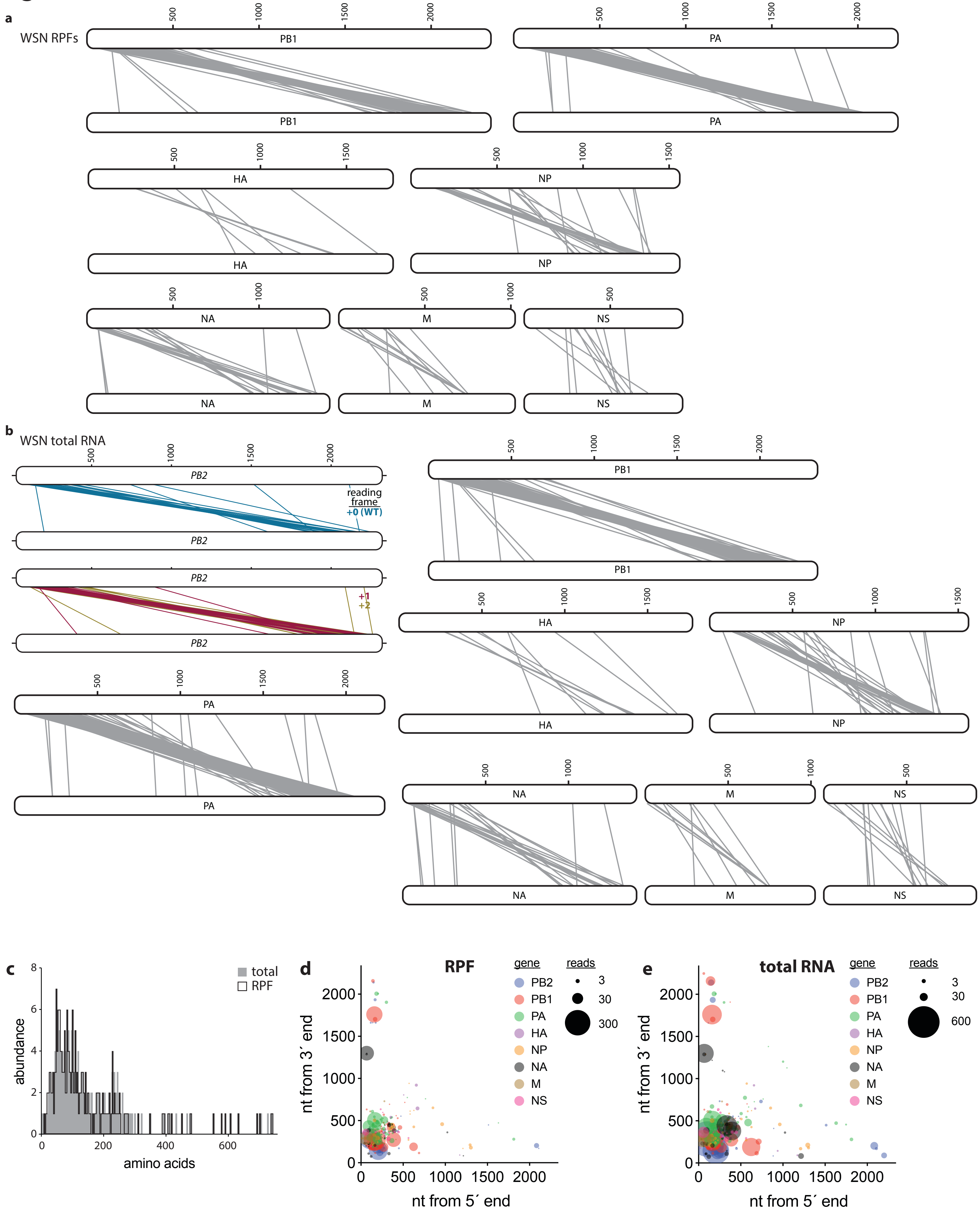
a
WSN RPFs
PB1
PA
HA
NP
NA
M
NS
500
1000
1500
2000
b
WSN total RNA
PB2
reading frame
+0 (WT)
+1
+2
PB1
PA
HA
NP
NA
M
NS
c
total
RPF
abundance
amino acids
d
RPF
gene
reads
PB2
PB1
PA
HA
NP
NA
M
NS
3
30
300
nt from 3´ end
nt from 5´ end
e
total RNA
600
