## Supplemental Figure 3 for "Cryptic proteins translated from deletion-containing viral genomes dramatically expand the influenza virus proteome"

**Figure S3: Parallel coordinates for DelVGs detected in human challenge studies. Associated with Fig 1.** (**a)** Humans were challenged with A/Wisconsin/67/2005 (H3N2) and DelVGs were identified by sequencing as described in Martin, et a. 2019. (doi:10.1101/814673). (**b)** DelVGs were identified by sequencing nasopharyngeal aspirates from patients naturally infected with A/Anhui/1/2013 (H7N9) as described by Lui, et al. 2019 (doi:10.1080/22221751.2019.1611346). For (a) and (b), polymerase genes are shown as plus-sense cRNA. Deletion junctions were remapped to full length versions of each gene. DelVG are depicted by a line connecting the 5´ (top) and 3´ (bottom) ends of each deletion. For PB2, the reading frame downstream of the deletion was predicted.

**Fig S3**

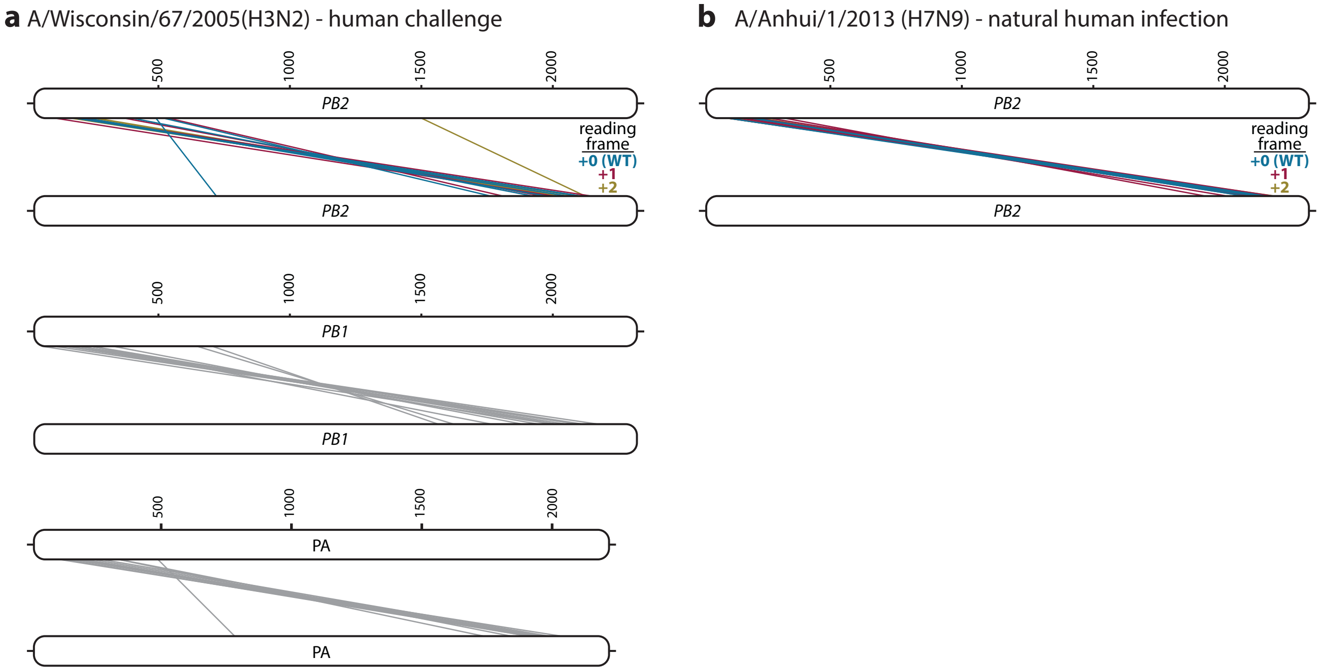
a A/Wisconsin/67/2005(H3N2) - human challenge
500
1000
1500
2000
PB2
reading frame
+0 (WT)
+1
+2
PB2
PB1
PB1
PA
PA
b A/Anhui/1/2013 (H7N9) - natural human infection
PB2
PB2
