## Supplemental Figure 4 for "Cryptic proteins translated from deletion-containing viral genomes dramatically expand the influenza virus proteome"

**Figure S4: Parallel coordinates for PB2 TRAP V5 reporter virus. Associated with Fig 3a-c.** DelVGs were identified by sequencing genomic RNA purified from virions of three plaque-purified stocks of the PB2 TRAP V5 reporter virus. **(a)** Data from each sample were consolidated and visualized by parallel coordinate mapping. Genes are shown as plus-sense cRNA. Each DelVG is depicted by a line connecting the 5´ (top) and 3´ (bottom) ends of each deletion. Coordinates can be found in Supplemental Table 4. **(b)** Junctions and read depth were determined for all DelVGs and plotted as a function of their positions from the ends of the respective viral gene. These are the same data as in Fig 3c, but scaled to show the full length of the gene segment. Note that none of the DelVGs in this sample mapped to *NS*. See also Supplemental Table 4 for underlying data.

### Fig S4

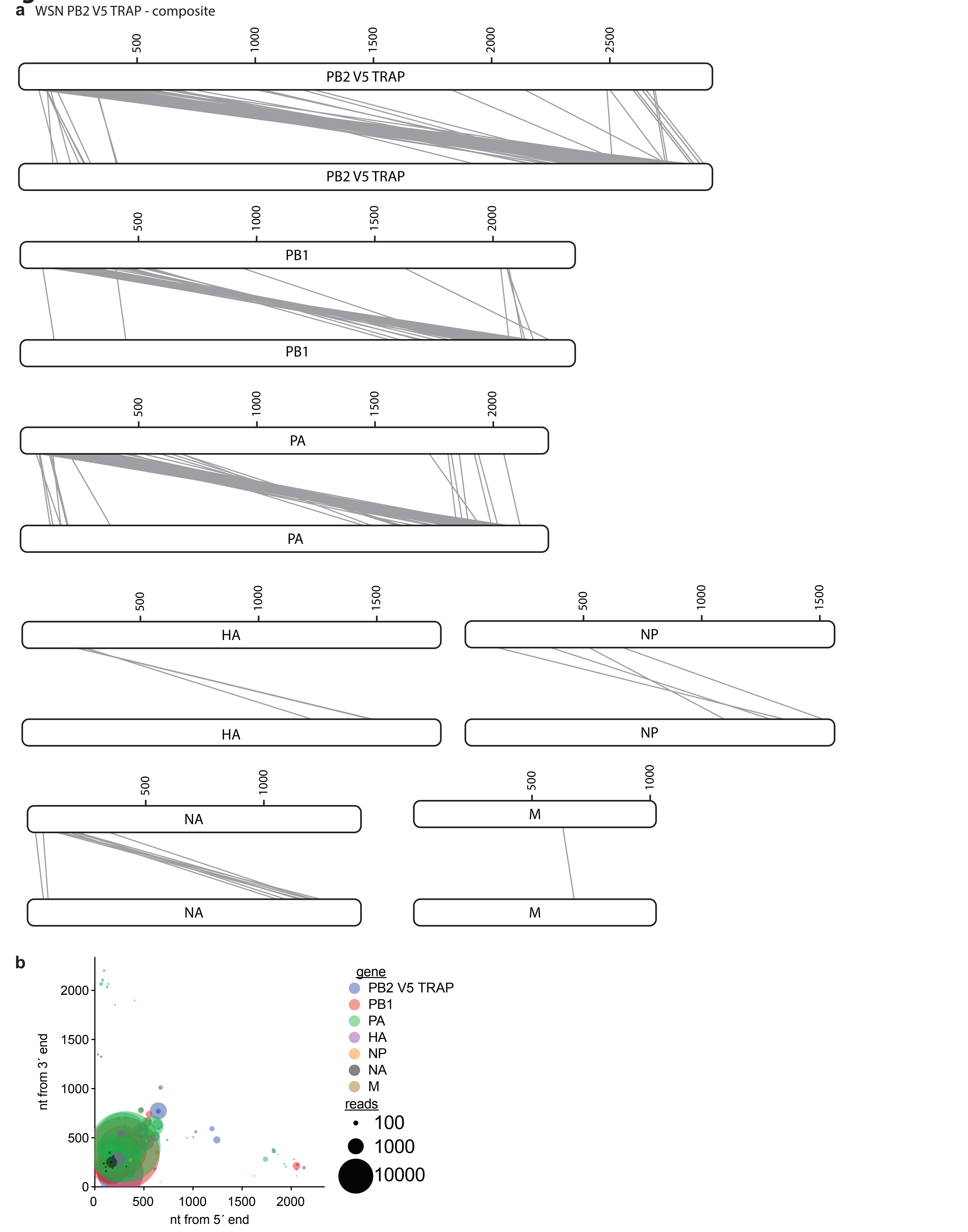
a
WSN PB2 V5 TRAP - composite
500
1000
1500
2000
2500
PB2 V5 TRAP
PB2 V5 TRAP
PB1
PB1
PA
PA
HA
HA
NP
NP
NA
NA
M
M
b
nt from 3´ end
nt from 5´ end
0
500
1000
1500
2000
gene
PB2 V5 TRAP
PB1
PA
HA
NP
NA
M
reads
100
1000
10000
