## Supplemental Figure 5 for "Cryptic proteins translated from deletion-containing viral genomes dramatically expand the influenza virus proteome"

### Figure S5: Uncropped blots from main figures

Fig 3D

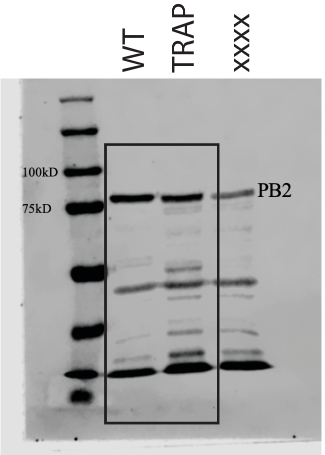


input
blotted for PB2

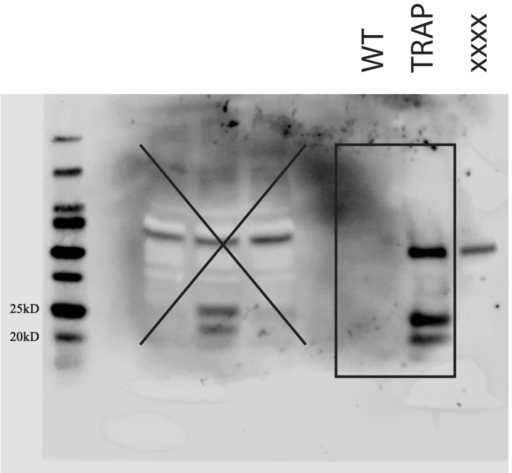


V5 IP
blotted for V5

Fig 4C

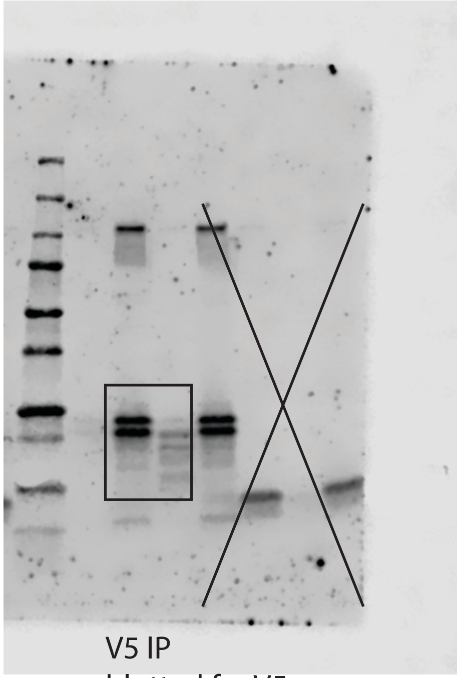

V5 IP
blotted for V5

Fig 4E

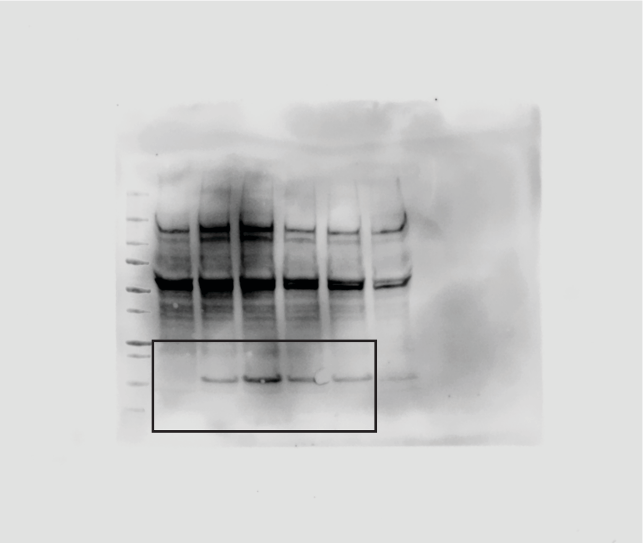

Fig 4F

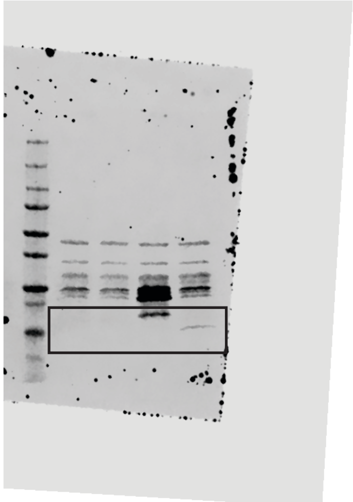

Fig 5

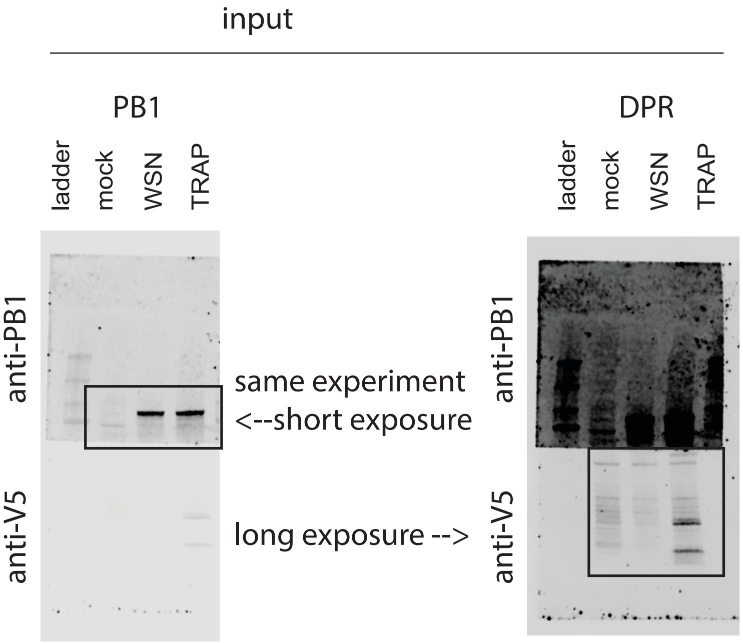


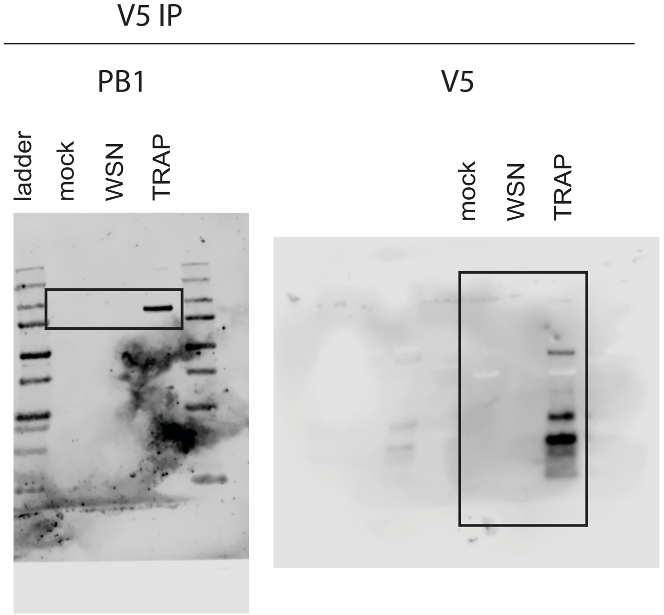


5B, left

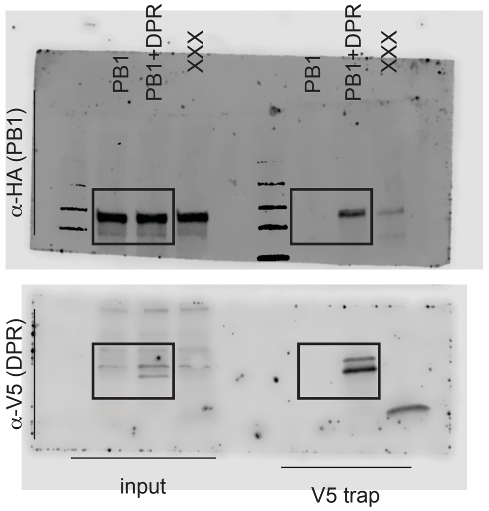


5B, right

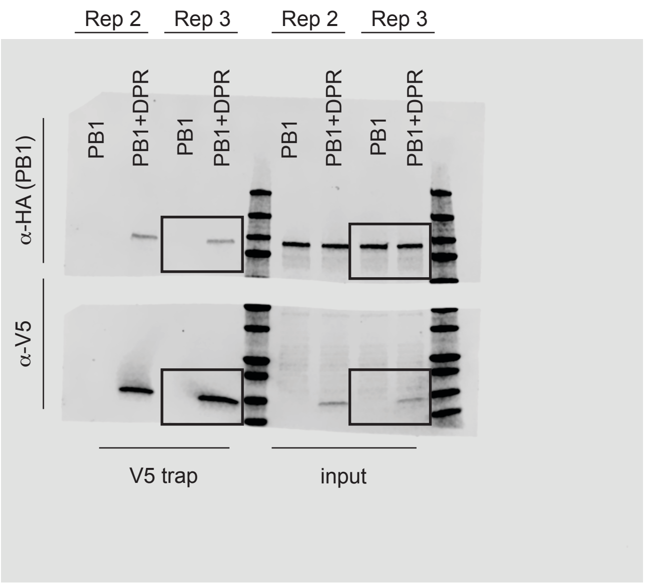

5C

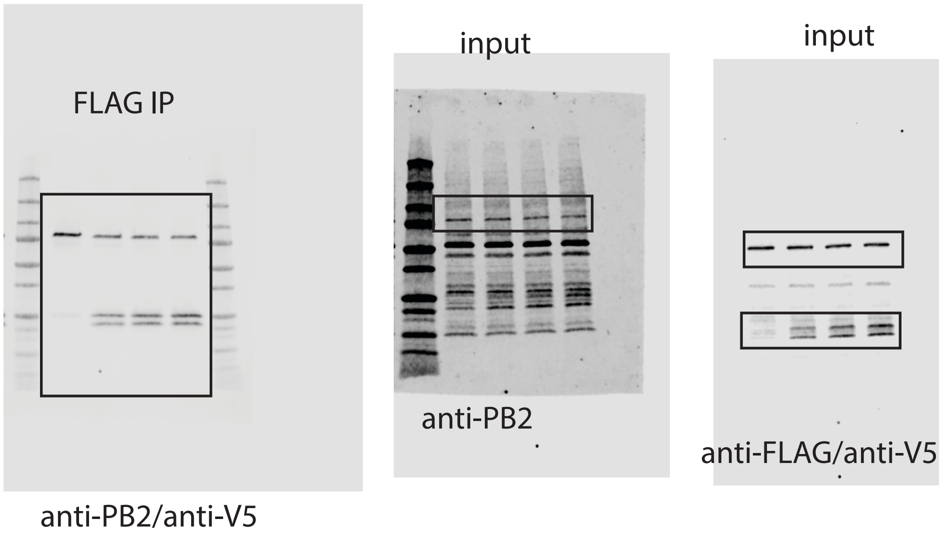
FLAG IP
anti-PB2/anti-V5
input
anti-PB2
input
anti-FLAG/anti-V5
